# Information synergy: adding unambiguous quality information rescues social information use in ants

**DOI:** 10.1101/219980

**Authors:** Tomer J. Czaczkes, John J. Beckwith, Anna-Lena Horsch

## Abstract

Animals have access to many alternative information sources when making decisions, such as private information (e.g. memory) and social information. Social insects make extensive use of social information. However, when intentional social information (e.g. pheromone trails in ants) conflicts with private information (e.g. route memories), insects often follow their private information. Why is this? We propose that an asymmetry in the type of information provided by these two information sources drives the neglect of social information: In ants, workers with certain information about the quality of a food source (memory) ignore valuable social information (pheromone trails) because the pheromone trails encode only a very ambiguous measure of food quality. This leads to a testable hypothesis: the addition of unambiguous quality information should rescue social information following. To test this, we trained ants to a poor quality (0.25M sucrose) food source, and then provided an alternative path along with either 1) no information, 2) a pheromone trail, 3) a 0.2μl 1.5M sucrose droplet, providing unambiguous quality information, or 4) both a trail and a droplet. When either no or only one information source was provided (1-3), most ants (60-75%) continued following their own memory. However, the addition of unambiguous quality information (4) rescued trail following: when both a trail and a droplet were provided, 75% of ants followed the trail. In further experiments, we show that quality information gleaned from direct contact with fed nestmates produced similar effects. Using florescence microscopy, we demonstrate that food (and information) flows from fed workers to outgoing foragers, explaining the frequent contacts on trails. We propose that the type of information an information source can convey, and its ambiguity, is a strong driver of which source of information is attended to.

## Introduction

Animals have access to many types of information, which come in various forms [1,2]. Two important classes of information are private information, which is available only to the individual animal, and social information, which is acquired from other animals. Private information includes genetic information and non-genetic information such as internal states and, importantly, memories. Socially-acquired information includes any information generated by the behaviour of another organism [2]. These may be social cues, such as the presence of conspecifics at a resource, or intentionally produced social signals, such as the waggle-dance of a honey bee.

Multiple information sources may provide information about the same thing. For example, both the colour and the odour of a flower patch, and whether other bees are dancing for that patch, might inform a bee about whether and where food is available in the environment [3,4]. These multiple information sources may interact in a variety of ways. Information may be used hierarchically, with one information type being used whenever it is available, and if it is not the next on the list is used [5]. When information sources agree, they may act additively or synergistically to improve behaviour. For example, ants which have private information about a food source (memory) which agrees with social information (a pheromone trail leading in the same direction) walk 25% faster than ants with either only one or none of these information sources [6]. These information sources also act additively by causing more ants to follow a pheromone trail [7]. Bumblebees which have experienced a novel flower odour in the nest (public information) are more likely to exploit the novel flowers, but only once they have tasted a flower scented with this novel odour (private information) [8].

A particularly interesting situation is when information sources conflict. One option in such situations is to weight information from different sources and produce an intermediate value [7,9,10]. An alternative is to rely on one type of information and ignore the others. In some situations, such as choosing a path at a trail bifurcation, an intermediate option is not available. Different strategies might be undertaken due to different ecological conditions or the state of individual animals, and understanding what strategies animals use to decide which information to follow is a very active field of research [2, 11–14]. Information use strategies can be broadly divided in to *when strategies* (when to use which information type) and *who strategies* (whose information to use) [15].

Evidence supporting various information use strategies has been described in wide range of taxa, including insects, fish, lizards, rats, and humans [16–19]. Information use strategies have been especially well researched in social insects such as ants and bees, as in these groups social information has the potential to play a very large role [12,20]. Social insects also represent a special case, since there are not expected to be any conflicts of interest in many situations requiring communication, such as foraging and nest-site selection. This should, at face value, strengthen the role of explicit social signals such as pheromone trails or waggle-dances. However, somewhat surprisingly, in most cases in which such intentionally produced social information is conflicted with private information, the ants and bees predominantly follow their own memories [21–29]. This was found not to be the case in only a few instances [23,30,31]. This preference could have strong implications for colony level behaviour, preventing colonies from optimally exploiting their environment.

Why do ants and bees so often underweigh or ignore the directional information provided by their nest mates? Explanations of information use strategies have been made in terms of the reliability of an information source – how often using one type of information is associated with a positive outcome or how outdated the information is (e.g. [14,32]), or the cost of acquiring new information [33–35]. However, information sources may provide information along multiple information dimensions. These different information types may each have a higher or lower information contents. For example, pointing in a direction can provide high precision and accuracy (high information content) in terms of direction but be ambiguous (low information content) in terms of distance. We hypothesised that information ambiguity along the quality dimension may be driving the neglect of social information. While pheromone trails provide highly accurate and precise directional information, they provide only very noisy, imprecise, and ambiguous information about the quality of a food source. Pheromone trail strength varies with resource quality, time since discovery, recruitment rates, and the individual pheromone depositions behaviour of ants: A strong trail may lead to a good food source exploited by a few ants, or to a poor one exploited by many ants; a weak trail may lead to a poor food source, or to a good one which has been unproductive for a while, or to a newly discovered food source. Importantly, there is a very large variation in the amount of pheromone deposited by individual ants to food sources of the same quality [36]. With no strong evidence that the new food source is better, foragers avoid paying the costs of attempting to find an advertised food source [33,37,38], and fall back on exploiting the food source she knows about.

This hypothesis leads to a testable prediction: If a worker foraging on a poor food source can be given unambiguous information that a better food source is available, she should follow the trail pheromone. This is because she should update the probabilities that following a pheromone trail will lead to a better food source than the one she is foraging on, improving the chances that the trail leads to better food. Here, we set out to test this prediction.

## Materials and methods

### Study species and maintenance

We used 8 queenless colony fragments of the black garden ant, *Lasius niger* (Linnaeus), collected from eight different colonies on the University of Regensburg campus. Colonies were housed in a plastic box (40×30×20cm) with a layer of plaster on the bottom. Each box contained a circular plaster nest (14cm diameter, 2cm high). Colonies contained c. 1000 workers and small amounts of brood. The ants were fed ad *libitum* on 1M sucrose solution supplemented with *Drosophila melanogaster* fruit flies. Colonies were deprived of food for four days prior to each trial to give high and consistent motivation for foraging and pheromone deposition. Water was provided *ad libitum*.

### Experimental series 1 procedure

10 different experimental treatments were carried out in this experimental series, coded A-J. An overview of the different treatments is provided in table S4 in the supplement. All treatments were variations on a central design. The aim of the experiments was to test whether ants with a well-established memory of finding food on one arm of a Y-maze can be induced to search a newly-presented alternate arm by the provision of various information sources. The information sources provided could be a pheromone trail, a small (0.2μl) droplet of sucrose (simulating trophallaxis or on-trail contact with a fed ant), a novel scent in either the droplet or on the newly presented path or both, or a combination of these. The scent treatments were added to test whether similar droplet and pathway odours cause a further increase in trail follow, as there could thus be an associative link between the quality of the droplet and the odour of the new runway [39]. However, as there was no effect of the various odour treatments (see supplement S1 for details) we pooled our data into treatments groups for the final analysis, depending on the sources of information available to the ants, and the quality of the original feeder to which it was trained. This resulted in 5 treatment groups: i) no information (treatments A and B), ii) only a droplet of 1.5M sucrose just before the bifurcation (treatments C-E), iii) only a pheromone trail on the new path (treatment F), vi) both a pheromone trail and a droplet (treatments G-I), and finally v) a pheromone and a droplet for ants trained to 1.5M sucrose (treatment J).

A trial began by allowing an ant onto the apparatus via a drawbridge. The drawbridge led to a Y-maze with one arm drawn out just out of reach of the ants, to form an L maze (Figure 1). The arm that was out of reach (left or right) was systematically varied. The arms of the maze were 10cm long and 1cm wide, narrowing to 2mm wide at the junction. The arms of the maze were covered with paper overlays. The overlay on the stem of the maze was unscented, while the overlay on the arm of the maze was scented with either rosemary or lemon essential oil – the odour used was systematically varied between trials. Odour impregnation was achieved by storing the paper overlays in a sealed plastic container containing one 0.1ml drop of essential oil on a glass petri dish for at least 24 hours. An acetate sheet affixed to the end of the L-maze arm acted as a feeder. A large drop of 0.25M sucrose solution (1.5M in treatment J), flavoured similarly to the L maze arm leading to it, was placed onto the feeder. Sucrose solutions were flavoured by adding 10μl essential oil per 100ml sucrose solution, following *Czaczkes et al.* [40]. Once the ant found the feeder it was marked with a dot of acrylic paint on the abdomen, and allowed to return to the nest. While the ant was in the nest, unloading her sucrose load, the paper overlays on the stem and arm were replaced by fresh overlays, to remove any trail pheromone the ant might have laid. The ant was then allowed to make 3 further return visits to the same feeder, resulting in 4 visits to the original feeder in total. This is sufficient to ensure that, even given the relatively low quality of food, most ants will return to this arm of the maze given a choice [41].

**Figure 1.**
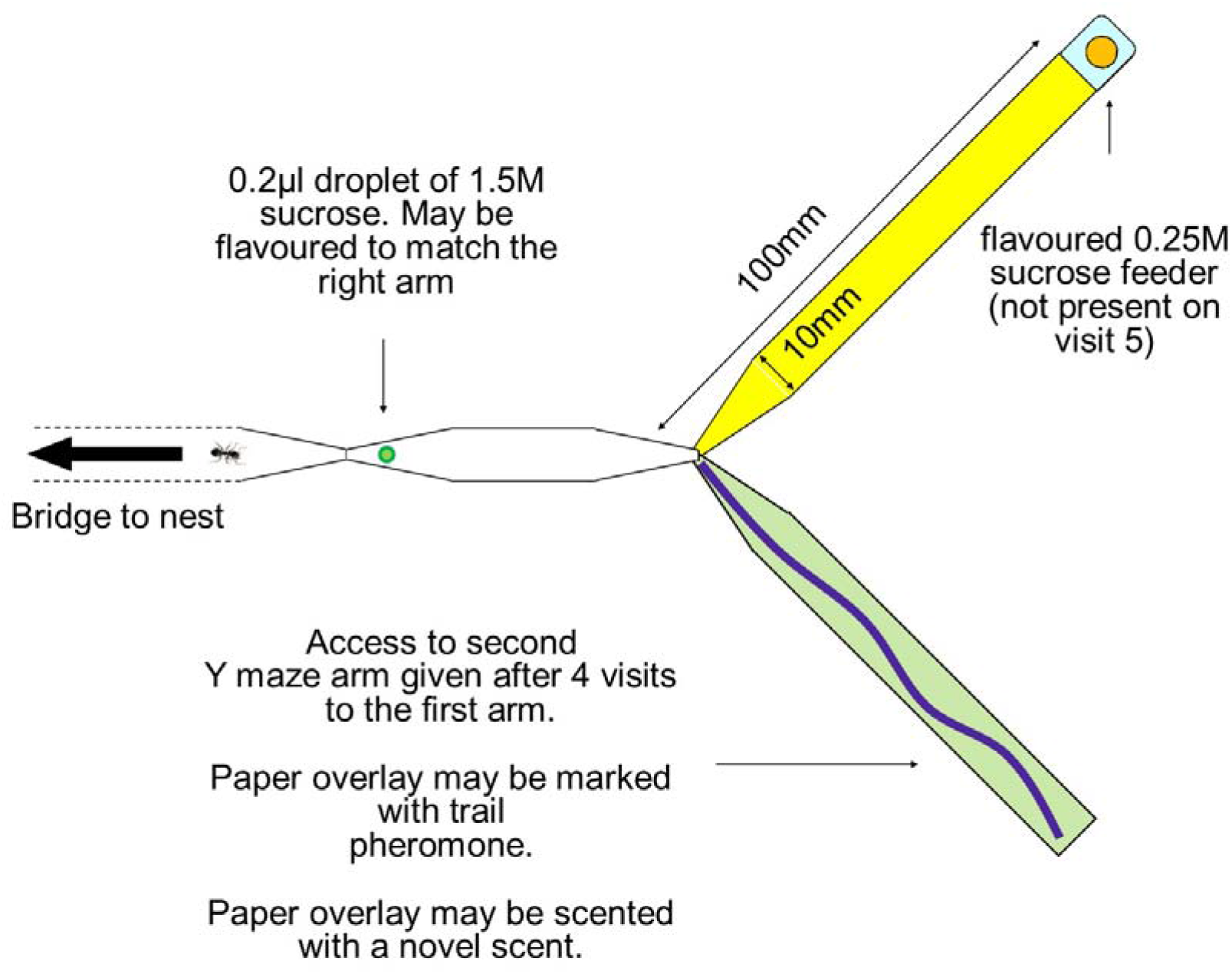
experimental setup. Ants are first trained for 4 visits on a Y-maze with access to only one arm (here the top arm, in yellow in the figure but not coloured in the experiment), leading to a 0.25M sucrose feeder flavoured with lemon or rosemary (1.5M in treatment J). On the 5 outward journey the second arm of the Y-maze (here bottom, green) is added, and the feeder removed. The new arm may be marked with a pheromone trail and/or scented with a novel scent (lemon or rosemary). Additionally, a small (0.2μl) droplet of 1.5M sucrose may be placed after a narrowing of the stem before the bifurcation, which the ant can drink but will not become satiated from. The droplet may be flavoured to match the new Y-maze arm scent or may be unflavoured.

While the ant was in the nest prior to its fifth return to the feeder, the feeder from the original arm of the Y-maze was removed, and access to the second arm was given. The new arm was either marked with a pheromone trail (treatments F-J) or not (treatments A-E). This pheromone trail was produced by immersing 8 worker hindgut glands in 2ml of dichloromethane (DCM), following von Thienen *et al.* [42]. 5.6μl of this mixture was applied in an even line along the paper overlay covering the arm, using a capillary tube (Servoprax GmbH, Germany). This amount was calculated to produce a pheromone trail of a realistic strength [42], and in control trials elicited a trail following accuracy of 82% (594 / 715), which is indistinguishable from those recorded from naïve ants following a reasonably strong naturally-deposited trails as reported by [43] (82-83% accuracy). See supplement S1 for detailed methods and results regarding the artificial pheromone trail.

The new arm was also either scented (treatments B, D, E, H, I, J) with a different scent to that of the original arm, or was not (treatments A, C, F, G). Lastly, a small (c. 0.2μl) droplet of sucrose either was (treatments C, D, E, G, H, I, J) or was not (treatments A, B, F) placed on the stem of the Y-maze, before the bifurcation but just after a narrowing of the stem. This ensured that the ant contacted the droplet as it walked towards the bifurcation. This droplet was either flavoured similarly to the scent on the newly presented runway (treatments E, I, J) or was unscented (treatments C, D, G, H). The droplet was large enough for the ant to detect and drink, but not enough to satiate the ant [44], which proceeded onwards after drinking the droplet. This droplet was designed to simulate trophallaxis or contacting fed ants on the trail. Trophallaxis is used in the nest to unload food, and also provides ants with information about the food available in the environment [45–48]. This information is attended to very strongly [46,49]. *L. niger* ants very often contact fed workers on their way to a food source, and can often be observed performing trophallaxis away from the nest. Recipients of such on-trail trophallactic interactions usually continue on their outwards journey afterwards (TJC, personal observation). Unfortunately, we could not reliably achieve such on-trail trophallaxis with the trained ant under controlled experimental conditions. However, we ran experimental series 2 (see below) to simulate on-trail contacts with other ants.

We then noted which arm of the Y-maze the ant chose. We took two choice measurements: the initial decision, as defined by the ant crossing a line 2cm from the bifurcation, and the final decision, as defined by the antennae of the ant reaching the end of the Y-maze arm. As the ant reached the end of the Y-maze, it was allowed to walk onto a piece of paper and replaced on the path leading to the Y-maze, before the location of the sugar droplet. This allowed us to make 10 repeated measures of each ant. After 10 such measurements the ant was permanently removed from the colony. The number of ants tested in each treatment is given in table S7 in the supplement.

### Experimental series 2 procedure

The aim of this experimental series was to test the main findings of series 1 (see figure 2) in a more biologically realistic way. Training and tests were identical to treatment G in series 1, i.e. ants were trained to find poor (0.25M) sucrose at the end of a Y-maze. On the 5 visit the unrewarded arm was marked with a pheromone trail. In this experiment the alternative Y-maze arm was present and accessible, but not rewarded, during training. Critically, unambiguous quality information was provided not via a drop of sucrose, but via nestmates. On all visits the outgoing focal ant was allowed to walk into a small (5×5cm) arena containing 5 nestmates. On the 4 training visits these nestmates had previously been fed to satiety on 0.25M sucrose. The focal ant was left in the arena for two minutes, and made on average 9.7 (SD 2.3) contacts with the fed ants. A contact was defined as the front half of the focal ant contacting the front half of any of the other ants. On several occasions (13% of visits, 47/357) trophallaxis occurred, and this was noted. After this contacting phase the ant was allowed to continue towards the Y-maze. Thus, training was identical for all ants. Then, in half of the 5 (test) visit, the 5 contacting ants were fed to satiety on 1.5M sucrose. In the other half of the trails on the 5 visit these ants were fed with 0.25M sucrose as during training, constituting a control. Thus, ants experienced one of two treatments: 0.25M sucrose from the contact ants during the final visit, or 1.5M sucrose from the contact ants on the final visit.

**Figure 2.**
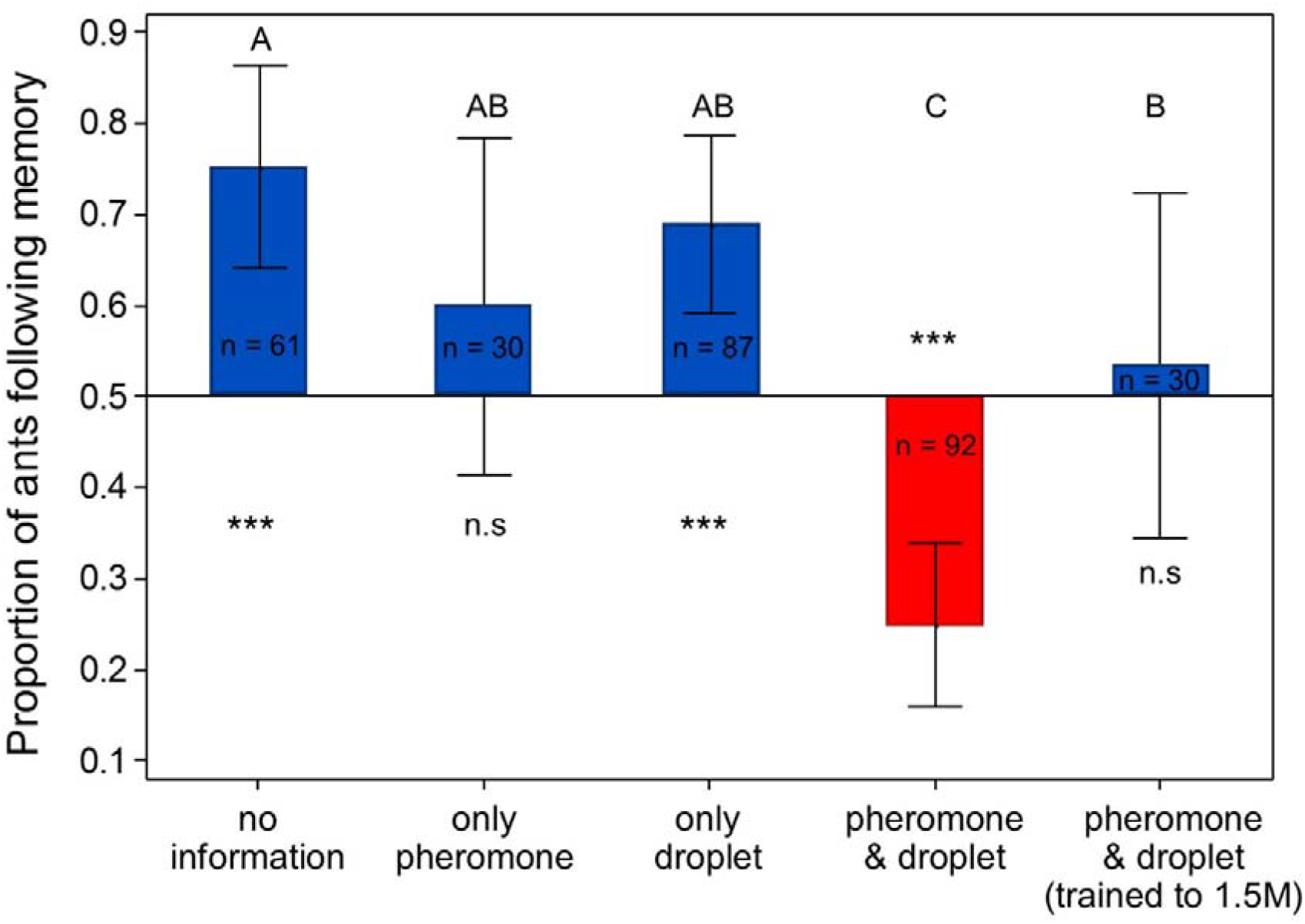
proportion of ants following their memory down the original Y-maze arm as a function of what other information was available to them. Ants may have had either no information about the new Y-maze arm, a pheromone trail on the new arm, a small 1.5M sucrose droplet before the bifurcation, or a pheromone trail and a droplet. In most groups ants were trained to 0.25M sucrose, but in one group they were trained to 1.5M sucrose. Whiskers are 95% C.I. for the mean. Letters indicate groups that are significantly (P < 0.005) different from each other. “***” and “n.s.” signify groups that are either significantly different from 0.5 (exact binomial test, P < 0.0001), or not different from 0.5 (P > 0.05), respectively. Sample sizes per group are show in the bars. Detailed statistical analysis results are presented in tables S1 and S2 in online supplement S1.

**Figure 3.**
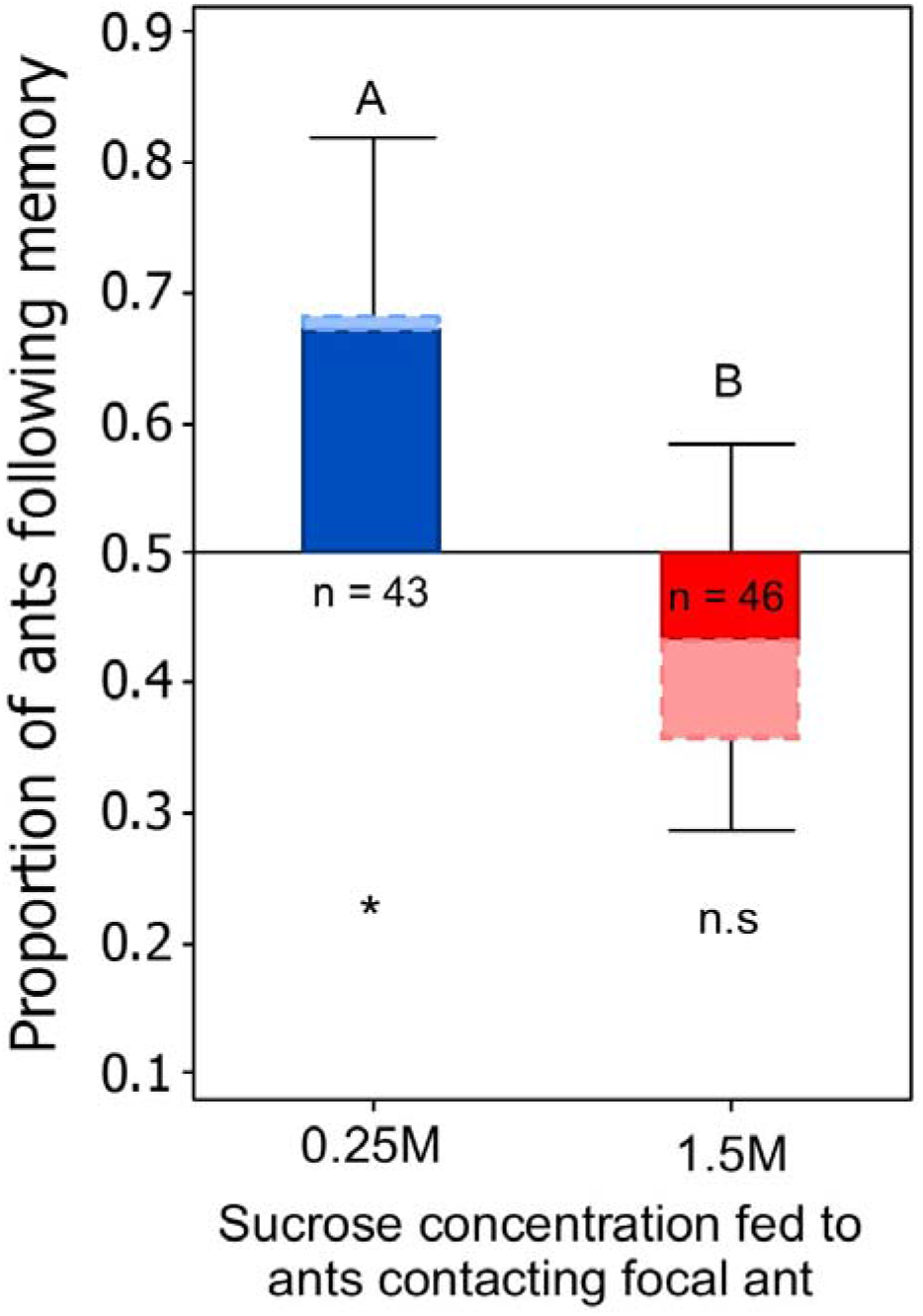
proportion of ants following their memory on a Y-maze rather than the alternate which was marked with trail pheromone. Ants encounter nestmates fed on either 0.25M (training quality) or 1.5M (higher quality) sucrose on the way to the choice point. Note that in c.30% of trials information transfer did not take place between the fed ants and the focal ant. The unadjusted proportion are given in the darker portion of the bars, and the 95% C.I. whiskers refer to these. The estimated adjustments are shown in the lighter portion of the bars, and are 0.69 for the 0.25M treatment, and 0.35 for the 1.5M treatment. These adjusted estimates are based on strong assumptions (see S1) and should be treated with caution.

#### Testing for information transfer during ant contacts

To test whether sugar solution, and thus information, is being transferred during these contacts, we used florescence microscopy to trace sugar flow between ants, as in [50]. A focal ant was allowed to find a 0.25M sucrose solution on one arm of a Y-maze and return to the nest, as above. On her second outward journey she was confined in a contact arena for two minutes with 5 nestmates that had been fed 1.5M sucrose containing 0.08 g/l] Rhodamine B. The focal ant was then removed, freeze killed, and the head, thorax, and abdomen separated and crushed between two microscope slides. The ant was then examined under green laser light (555nm) using a Zeiss Axioplan-2 fluorescence microscope for presence of Rhodamine B. Control ants taken directly from the nest were also examined.

### Statistical analysis

Since the initial decision (crossing a line 2cm from the bifurcation) and the final decision (reaching the end of the arm) rarely differed (<5% of cases), we analysed only the final decision. While we collected 10 data points per ant to increase the potential power of our experiments, we finally decided to perform a conservative analysis by only considering the first decision made by the ants. This avoids the possibility of ants becoming frustrated in later trials. Initial data exploration revealed a leftward side bias, which is not uncommon in the behaviour of ants or other animals [51], and so training side was also added to the model. Finally, colony identity was added as a random effect. This resulted in the following model:

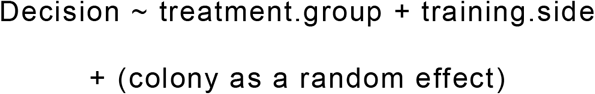

As the data is binomial we used a binomial error structure. Data were analysed in R v.3.4.1 [52] using generalised linear mixed models from the LME4 package [53]. Model fit was checked using the DHARMa package [54].

We also tested whether the decisions of each group differed significantly from random using two tailed exact binomial tests. Due to the limitations of using p-values alone, we provide estimates and confidence intervals for all effects.

## Results

The type of information available to a forager strongly influenced her decision of whether or not to follow her memory. For ants trained to 0.25M sucrose, given just an unmarked novel path on the Y-maze, most ants (75%) choose to follow their own memory (figure 2), significantly more than chance (two tailed binomial exact test, 46/61 followed their memory, mean = 0.75, 95% C.I. range = 0.63 – 0.86, p < 0.0001). The addition of a small 1.5M sucrose droplet does not significantly change the choices made by the ants, who still preferentially follow their memory (binomial test, 60/87, mean = 0.69, 95% C.I. = 0.58 – 0.78, P = 0.0005). The addition of a pheromone trail on the novel arm still resulted in most ants (60%) following their memory, but this result was not different from random choice (binomial test, 18/30, mean = 0.6, 95% C.I. = 0.41 – 0.77, p = 0.36). These three groups do not differ significantly from each other (GLMM, z ≤ 1.90, p ≥ 0.057, for details of all pairwise comparisons, including estimates for each comparison, see table S1 in supplement S1).

However, a striking synergistic effect of presenting a pheromone trail and a sucrose droplet was found. If both were presented, only 25% of ants chose to follow their memory. This is significantly fewer than chance (binomial test, 23/92, mean = 0.25, 95% C.I. = 0.17 – 0.35, p < 0.0001) and differs significantly from all other groups provided with less information (GLMM, Z ≥ 2.97, P ≤ 0.0029).

The effect of presenting both information sources depends on the relative quality of the droplet presented and the food quality the ant was trained on. If ants were trained on 1.5M sucrose, 53% of ants chose to follow their memory, which does not differ from random choice (binomial test, 16/30, mean = 0.53, 95% C.I. = 0.34-0.72, p = 0.86). This is significantly greater than for ants receiving the same information, but trained on poorer food (GLMM, z = 0.23, p = 0.020), and also significantly less than the ants receiving no information (GLMM, z = −2.56, p = 0.011). It is not significantly different from the ants receiving either one information source (GLMM vs only droplet, z = 1.95, p = 0.051, vs only pheromone z = 0.60, p = 0.55).

Similar results were found from experimental series 2, in which unambiguous information about available food quality was provided via contact with fed nestmates. The behaviour of focal ants which contacted ants fed on either 0.25M was significantly different from those contacting 1.5M-fed ants (GLMM, z = 2.31, P = 0.021). While focal ants which contacted 0.25M-fed nestmates preferentially followed their memory to the original food side (binomial test, 29/44, mean = 0.66, 95% C.I. 0.50-0.80 p = 0.049), focal ants which contacted nestmates fed with 1.5M tended to follow the pheromone trail, although their choice was not significantly different from chance (20/47, mean 0.43, 95% C.I. 0.28 – 0.58, p = 0.38). However, it is likely that not all focal ants received information from the fed nestmates, making the reported 43% an overestimate. In the fluorescent sucrose test for information transfer, only 70% of focal ants received marked food from their fed nestmates. Thus, we should expect around 30% of our test subjects in experiment series 2 not to have received unambiguous food quality information, and thus follow their memory at a rate of around 60% (see figure 2 ‘only pheromone’). Adjusting for this rate of information transfer failure, we estimate 68.7% of ants in the 0.25M treatment followed their memory, compared to 35.1% in the 1.5M treatment (see S1 for our calculations).

The entire dataset on which these results are based in provided in online supplement S2. All fluorescence images are provided in S3.

## Discussion

### Information sources and information conflict

Animals have access to many sources of information, which come in various forms [1,2]. Two important classes of information are private information, which is available only to the individual animal, and social information, which is acquired from other animals. When these information sources conflict, social insects often follow their private information [21–29]. We found that ant foragers preferentially follow their private memory over a pheromone trail if they have no further information about the quality of other potential food sources (figure 1a), or if this information implies the same low quality (figure 1b). However, if outgoing foragers are informed about a better potential food source, either via finding a small drop of better food or contacting nestmates which have fed on such food, they strongly prefer to follow social pheromone trail information. Why is this, and what can this tell us about the information use strategies animals employ?

### Information type is a key driver of information use decisions

Information use strategies have been divided into ‘when’ and ‘who’ strategies – when should information be used, and who should one listen to [12,15]. Here, we propose considering information use in terms of a ‘*what*’ strategy – what type of information does an information source convey? The dimensions along which information sources can be informative – the type of ‘data’ an information source conveys - is extremely relevant to deciding how to act on information. Information sources can provide multiple dimensions of information, each with a different level of precision, accuracy, and ambiguity. For example, a pointing finger can provide high precision and accuracy in terms of direction but be ambiguous in terms of distance. Different information sources may take precedent, depending on which information dimension is most important. Pheromone trails and honey bee waggle-dances do, in fact, encode information about food quality: individual foragers only recruit to sufficiently high quality food sources [55]. However, this information is extremely imprecise and noisy both due to individual variation in responses to fixed food quality [36,56] and, for pheromone trails, due to being ‘overwritten’ by other ants or evaporating. Private sensory information, and contacts with nestmates which provide this, *does* provide unambiguous quality information. We propose that it is this ambiguity along the quality dimension in some social information, or the certainty of the private information, which causes otherwise valuable social information to be neglected. Our data supports this claim. Explanations of information use strategies have been made in terms of the reliability of an information source – how often using one type of information is associated with a positive outcome, or how outdated the information is (e.g. [14,32]), or the cost of acquiring new information [33–35]. However, the type of information provided by an information source is often not explicitly considered. Some information sources, such as the waggle dance or a pheromone trail, provide unambiguous and precise information along one dimension, but ambiguous information along another. The neglect of ambiguous options, and the preference for certainty, are robust economic behaviours in humans, and have also been demonstrated in non-human primates [57,58,58,59].

### Information type asymmetry explains information use patterns described in other studies

Understanding information use strategies in terms of information asymmetry can explain previously puzzling findings. Leadbeater and Florent [8] unexpectedly found that bumblebees do not rate social information above personal experience, even when their personal experience becomes outdated. Bumblebees which had experienced a novel odour in the nest (social information) were just as unlikely to sample novel flowers scented with this odour as bees which did not experience the novel odour in the nest. Both groups continued choosing known-odour flowers, even when those flowers became unproductive. However, once the bees with the social information did, eventually, sample the novel flowers, they were much more likely to switch to the novel flowers than the group without social information. This is directly analogous to the situation we report: neither private sensory information providing reliable information in the dimension of quality (sampling the novel flowers), nor social information (smelling the novel odour in the nest) is sufficient to make bees change their choices. However, when both are provided together (here, after the first sampling of the novel flower), bees are willing to switch.

Similarly, understanding information use in terms of information quality can explain situations in which social information does override personal information. This is because in such cases, the social information does provide unambiguous quality information. For example, Dunlap et al. [14] and Smolla et al. [60] find that social information alone can drive decision making during flower choice in bumblebees, and that social information even outweighs private information when the two compete [14]. This result could be understood through the psychological concept of blocking [20]. However, differences in information content between the two information sources can also explain these results: in these studies the social information is based on making associations between a social cue (the presence of a conspecific) and the direct personal experience of the forager with the flower and its’ quality [20]. There is thus no quality ambiguity. Indeed, if ambiguity exists, it is lower in the social information: the forager not only has her own experience of finding the flower acceptable, but also the presence of a conspecific suggests that conspecifics also find the flower acceptable, resulting in more weight of information about quality for the socially advertised flower. Likewise, Josens et al. [49] describe how trophallactic interactions with a forager providing scented, untainted food can cause foragers to accept similarly scented food tainted with poison, which they would otherwise avoid. Here again, the social information component is based on direct sensory experience of the food (trophallaxis), and the reasons the ants overweigh this information over their own later sensory information are similar to those described for the bumblebee examples above.

### Direct transmission of quality information to foragers enables strategic information use

Outgoing ants repeatedly encounter returning ants on a trail. We have shown that during these contacts, small amounts of food, and thus information, can be transferred to the outgoing ant. Information transfer can be either via trophallaxis, or by detecting food remains smeared on the fed ant while it was feeding. We also observed transfer of food *to* active foragers from ants inside the nest, as reported recently in another species [50]. Similarly, dancing honeybees often engage in trophallaxis with followers. Ants receiving trophallaxis may even respond as if they have themselves successfully foraged, depositing trail pheromone when otherwise only successful ants do so [61]. Information gained during such contacts, and especially trophallactic interactions, seems to be particularly well attended to – even more so than information gained while foraging [46,49]. This information, gained in the nest or on the trail, can then drive strategic information use. An outgoing forager encountering returning ants which collected poorer food may continue exploiting her own known food source, while outgoing foragers encountering more successful returning ants may begin to follow pheromone trails, thus playing a “copy if better” strategy. The “copy if better” strategy is uncommon in biology, as direct quality comparison is not usually possible and accurate social comparison may require advanced cognitive abilities [62]. If a forager has only found poor quality food, and encounters only other foragers with equally poor food, it may be more likely to begin scouting, following an “innovate if dissatisfied” strategy. Thus, provision of unambiguous quality information may well be a major role of interaction of ants on a trail. Note that while in these experiments only one alternative path is offered, in nature many alternative paths will be available, some marked with pheromone and others not. Thus, the unambiguous quality information provided by contacts in the nest and on the trail can drive general information use strategies, but do not specify which precise path to take. It is thus all the more striking that such information provision causes a reversal in information use choice, as the quality information is not directly linked to the alternative path. Provision of information about food associated cues, such as the odour of the shared food [45,49], is also no doubt an important role of contacts and trophallaxis, although we could not find any role for food odour in the current experiment (see supplement S1 figure S1). The ability to redirect workers to the highest quality underexploited food source will aid colonies secure resources from competition [63] and most efficiently utilize a colony’s’ limited processing and storage capacity.

### ‘Inactive’ workers in a colony may synthesise and disseminate foraging information

Trophallactic networks within a nest are complex [50,64]. Social insect colonies contain a high proportion of apparently taskless individuals ([65] and references therein), which can be divided into “inactives” and “interactives” [66]. Interactives have a more central place in interaction networks within the colony [65,66]. If they also receive trophallactic interactions from many foragers and food receivers, they may act to homogenise the various food qualities being returned to the nest, providing a reading of the average food being brought into the colony. Foragers interacting with these ants may then compare this reading to their current exploited food source. If they find their own food source to be worse, they may begin following social information. Using newly developed florescent imaging technologies allows the flow of food via trophallaxis in a colony to be tracked and quantified [50], opening the possibility to empirically test these suggestions. In line with this reasoning, ants which receive high quality food via trophallaxis are less willing to accept medium quality food, and vice versa for ants receiving poor quality food [67]. It is also known that ants which have experienced higher quality food source avoid exploiting poorer but otherwise acceptable food source discovered en-route [68]. It seems likely that ant foragers are very well informed about the various food qualities available in their environment, and scout, follow pheromone trails, or exploit known food sources adaptively depending on this information.

### Summary

Here, we explored how three different information sources, one private and two public, interacted. In doing so, we found evidence that the type of information an information source conveys is a key driver in whether it will be heeded or not. Some information sources may be highly informative about certain things, but nonetheless be neglected if they are uninformative along a critical informational dimension. We propose that as well as considering ‘when’ and ‘who’ strategies for information use, ‘what’ is an important question to ask: what type of information can be provided. Social insects, and animals in general, demonstrably use complex mechanisms to decide which information to attend to, and integrate information from a wide variety of sources in order to do so.

## Acknowledgements

Thanks to Wolfhard von Thienen for advice on producing an artificial pheromone trail of a realistic strength, to Stephan Schneuwly for the use of the florescent microscope, to Ofer Feinermann, Efrat Greenwald, and Lior Baltisansky for ideas relating to experimental series 2 and for providing Rhodamin B, to Ellouise Leadbeater and Christoph Grüter for comments on previous versions of this manuscript.

